# Position-Specific Enrichment Ratio Matrix scores predict antibody variant properties from deep sequencing data

**DOI:** 10.1101/2023.07.10.548448

**Authors:** Matthew D. Smith, Marshall A. Case, Emily K. Makowski, Peter M. Tessier

## Abstract

**Motivation:** Deep sequencing of antibody and related protein libraries after phage or yeast-surface display sorting is widely used to identify variants with increased affinity, specificity and/or improvements in key biophysical properties. Conventional approaches for identifying optimal variants typically use the frequencies of observation in enriched libraries or the corresponding enrichment ratios. However, these approaches disregard the vast majority of deep sequencing data and often fail to identify the best variants in the libraries.

**Results:** Here, we present a method, Position-Specific Enrichment Ratio Matrix (PSERM) scoring, that uses entire deep sequencing datasets from pre- and post-selections to score each observed protein variant. The PSERM scores are the sum of the site-specific enrichment ratios observed at each mutated position. We find that PSERM scores are much more reproducible and correlate more strongly with experimentally measured properties than frequencies or enrichment ratios, including for multiple antibody properties (affinity and non-specific binding) for a clinical-stage antibody (emibetuzumab). We expect that this method will be broadly applicable to diverse protein engineering campaigns.

**Availability:** All deep sequencing datasets and code to do the analyses presented within are available via GitHub.

**Contact:** Peter Tessier

**Supplementary information:** Supplementary data are available at Bioinformatics online.

## 1. INTRODUCTION

Protein therapeutics are a dominant class of therapeutic agents being used to treat myriad disorders and diseases. Their enormous design space enables the generation of diverse bioactive molecules, including hormones, enzymes, and monoclonal antibodies. In particular, antibodies are attractive therapeutic agents due to their high target affinity, low off-target binding, ease of production, and favorable biophysical properties (Tiller *et al*. 2015; Jain *et al*. 2017). Antibodies have been used to treat several different disease groups, including cancer, autoimmune disorders, and neurological diseases, leading to >100 approved antibody drugs and many more currently in clinical development (Lu *et al*. 2020; Carter and Rajpal 2022; Lyu *et al*. 2022; Kaplon *et al*. 2023).

The discovery of lead antibody candidates is commonly performed using methods such as animal immunization or *in vitro* antibody library screening techniques such as phage or yeast-surface display. The use of *in vitro* library screening enables the sorting and deep sequencing of large antibody mutant libraries to discover and optimize potent therapeutics. Indeed, recent advances in next-generation sequencing (NGS) has made it routine to sequence antibody libraries to select clones with high affinity and specificity (Glanville *et al*. 2015; Wrenbeck *et al*. 2017; Rouet *et al*. 2018).

However, it remains surprisingly challenging to identify universal metrics for selecting optimal antibody variants from deep sequenced libraries. Several simple metrics have been used in the past such as frequency, or the abundance of an antibody sequence found after selection, which assumes that variants with the highest affinities become most prevalent during the sorting process (Ravn *et al*. 2010, 2013; D’Angelo *et al*. 2014; Hu *et al*. 2015; Lopez *et al*. 2017; Barreto *et al*. 2019; Ferrara *et al*. 2020). Another common metric used to select antibody variants is the enrichment ratio, which is the frequency of the variant in the output of the selection divided by the frequency of the same variant in the original library (Maranhão *et al*. 2020; Kelil *et al*. 2021). However, both methods are limited because they disregard the vast majority of the information in the deep sequencing datasets and solely rely on the frequencies of each antibody variant of interest. Further, the uncertainties in the corresponding frequencies and enrichment ratios, especially for rare antibody variants, can result in limited correlations between either deep sequencing metric (frequency or enrichment ratio) and antibody properties of interest (e.g., affinity) (Fowler *et al*. 2011; Kowalsky *et al*. 2015; Rubin *et al*. 2017).

A second general approach for selecting optimized antibody variant from large sequencing datasets involves training machine learning models on the enriched antibody sequences (Magar *et al*. 2021; Narayanan *et al*. 2021; Saka *et al*. 2021; Hanning *et al*. 2022; Makowski *et al*. 2022a; Wang *et al*. 2022). These techniques utilize large datasets to learn underlying patterns within the sequences to make predictions of optimal variants. Despite the great potential of this general approach, the main common limitations are 1) the required user expertise needed to train the models, 2) model-specific differences in assumptions and predictions, 3) the potential for overfitting the training data, and 4) the lack of model interpretability in some cases.

Here we have sought to combine the strengths of conventional (frequency based) analysis with methods that use entire deep sequencing datasets to develop a hybrid approach that addresses previous limitations. One well established method uses position-specific scoring matrices (PSSMs) for analyzing the relative frequency of each amino acid at each site within protein sequence datasets (Henikoff *et al*. 1994; Tatusov *et al*. 1994). These PSSMs can be used to score sequences to obtain a metric for the similarity of the sequence to the observed dataset. We reasoned that an analogous type of matrix, a position-specific enrichment ratio matrix (PSERM), could be developed that reported the site-specific enrichment ratio of each amino acid at each mutated site and a similar scoring method would yield a composite score correlated with each antibody property of interest. Herein, we demonstrate the utility of this approach and how it compares to conventional methods for optimizing antibody affinity and non-specific binding.

## 2. METHODS

### 2.1 Datasets

In this paper, antibody deep sequencing data from three different selection campaigns were used that are briefly described below and summarized in **Table S1. Project 1** contains data from a recently published report aiming to decrease the non-specific binding of a clinical-stage antibody, emibetuzumab, while maintaining affinity to the antigen (Makowski *et al*. 2022b). This dataset contains sequencing data from the input library, positive selections for two different concentrations of antigen (0.1 and 1 nM hepatocyte growth factor receptor, HGFR), positive and negative selections for ovalbumin binding, and positive and negative selections for binding to a polyspecificity reagent (soluble membrane proteins). This dataset also includes the measured binding to both antigen and ovalbumin for a set of 125 antibodies. **Project 2** contains data from another antibody selection campaign aimed at improving the biophysical properties, such as self-association and host-cell protein binding, while maintaining high antigen affinity for a preclinical antibody specific for platelet derived growth factor BB. This dataset contains sequencing data from the input library, positive and negative selections against antigen, host-cell enzymes (LPLA2) and immunoconjugates (lenzilumab conjugated to quantum dots), the latter of which has been used as a surrogate for evaluating antibody self-association (Makowski *et al*. 2022a). Finally, **Project 3** contains data from a selection campaign for affinity maturation of an anti-Aβ antibody (Desai *et al*. 2021). This dataset contains sequencing data from several successive rounds of positive selection against Aβ fibrils, namely rounds 2, 3, 5, 6, and 7. For this dataset, the round 2 sample was used as the input sample and rounds 3-7 were used as the output samples. Details regarding the library sorting conditions are given in **Table S2**.

### 2.2 Deep sequencing of sorted antibody libraries

Deep sequencing datasets were prepared in the same manor explained elsewhere (Desai *et al*. 2021), unless noted otherwise. Briefly, sorted yeast-surface display libraries were yeast mini-prepped (Zymo, D2004) according to the manufacturer’s protocol. The antibody variable domain of interest was amplified from the isolated yeast plasmids with primers that added Illumina p5 and p7 adapter regions as well as i5 and i7 indexes to allow for demultiplexing. The primers were also designed to incorporate phasing of the PCR products to improve sequencing performance (Wu *et al*. 2015). All PCR products were gel purified on 1% agarose gel. The concentrations of the prepared DNA samples were then measured with a Qubit 4 1x High sensitivity analysis kit (Thermo Fisher, Q33231) and pooled in equimolar concentrations. The pooled DNA samples were then submitted to the University of Michigan Advanced Genomics Core for 2×300 bp paired-end MiSeq runs. All library samples were deep sequenced twice, and a summary of the number of reads per sample, number of unique sequences for both replicates, and theoretical library sizes are provided in **Table S3**.

### 2.3 Analysis of fastq files

To generate paired sequence-frequency datasets to use for analysis, the output fastq files from the sequencer were analyzed in several steps. First, the two reads were merged using BBMerge (**Project 1** and **2**) or FLASH (**project 3**) (Magoč *et al*. 2011; Bushnell *et al*. 2017). The file containing the correctly merged sequences was converted into a fasta file. The following quality criteria were imposed for a sequence to be included in the final dataset: i) the sequence had to be of the correct length; ii) it had to have the correct starting sequence; and iii) lack ‘N’s in the DNA sequence. The DNA sequence was then translated using Biopython (Cock *et al*. 2009). Because MiSeq has an error rate of approximately 1 error in every 1000 bases read, this full-length protein sequence was trimmed to only include the amino acids present at the intended mutated positions (Stoler *et al*. 2021). This sequence was termed the mutation string, and all sequences were represented by their associated mutation string. The number of times each residue string was observed was counted and this was converted into a frequency by dividing by the total number of observed sequences.

A matrix containing the average frequency of each clone in each sample was generated by merging the data from the previous step. Next, the dataset was trimmed to remove sequences that were not possible based on the library design, which resulted in removal of relatively few sequences. The counts of each clone in each replicate were added together and renormalized by the total number of counts in both replicates:

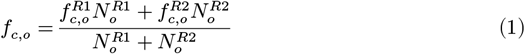

where 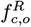 represents the frequency of clone *c* in sample *o* from replicate *R*, and 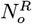 represents the total number of counts of all sequences in sample *o* from replicate *R*. All subsequent calculations were conducted based on this averaged frequency *f*_*c,o*_.

### 2.4 Creation of PSERM

The clonal frequency matrix was converted to a site-specific amino acid frequency matrix for each sample. Each observed sequence of a given sample, *i*, was iterated through and the counts of each amino acid at each position of that sequence were increased by the number of times that sequence was observed. After iterating through all the observed sequences, a pseudocount (defined as *s* and in this work set equal to 1) multiplied by a background probability (denoted as *p*_*a*_ and set equal to 1/20) was added to every amino acid at each position. The pseudocount was added to prevent problems when taking logarithms. Finally, this matrix was normalized by dividing by the total number of sequences, plus pseudocounts, in the sample. These matrices were mathematically defined as:

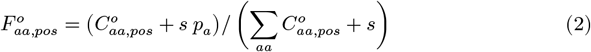

where 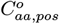 represents the count of amino acid *aa* at position *pos* in sample *o*. This matrix, and all other matrices in this work, were indexed by the amino acids, *aa*, for the rows and the positions, *pos*, for the columns. This frequency matrix, 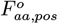, was used to construct a Position-Specific Scoring Matrix (PSSM) (Tatusov *et al*. 1994) by computing the log transform of the ratio of the matrix with the background probability defined as:

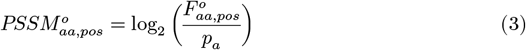

The frequency matrix of a given output sample, *o*, and the input frequency matrix, 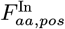, were also used to compute a **P**osition-**S**pecific **E**nrichment **R**atio **M**atrix (PSERM), which was the log_2_ transform of the ratio of the output sample frequency matrix to the input frequency matrix:

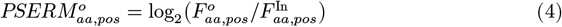

### 2.5 Scoring antibody sequences

In this work, antibody variants were scored in four main ways, namely in terms of their frequency, enrichment ratio, PSSM score, and PSERM score. The frequency of a clone was defined to be the averaged frequency of a clone in each sample according to **equation 1**. The enrichment ratio for a clone was defined as the log_2_ ratio of its output frequency to its input frequency:

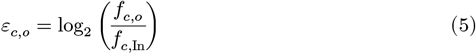

where *ε*_*c,o*_ denotes the enrichment ratio of a clone *c* for output sample *o*. In this equation, *f*_*c,o*_ was the frequency of a clone *c* in output sample *o*, and *f*_*c*,In_ is the frequency of the same clone in the input sample. The uncertainty in an enrichment ratio was calculated as previously reported (Kowalsky *et al*. 2015; Otwinowski *et al*. 2018) using the following equation:

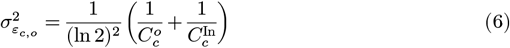

where 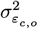 represents the variance of the enrichment ratio of a clone *c* in output sample *o* computed from propagation of error of the counts of the clone in the output sample *o*, 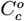, and the input sample, 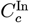.

Sequences scored with the PSSM or PSERM were scored by summing the values of their respective matrices according to the given sequence compositions. The PSERM scoring formula for a sequence *c* was given as:

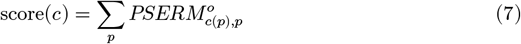

where *p* represents a mutated position and *c*(*p*) represents the amino acid of the scored clone at position *p*. The uncertainty in a PSERM score of a clone *c* can be computed using propagation of error assuming the counts of amino acids follow a Poisson distribution, as observed for the counts of individual clones. The uncertainty was given by the following equation:

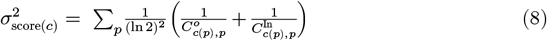

where 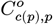 represents the count of the amino acid, *c*(*p*), of the clone *c* in position *p* in the output sample *o*, and 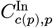 represents the count of the same amino acid in the input sample.

### 2.6 Epistatic effects

Epistatic effects, also known as epistatic shifts, of every residue sampled in the library were computed by comparing the enrichment profiles of every residue at every non-fixed position given a fixed residue at a fixed position. To do this, a fixed residue PSERM was computed and defined as a PSERM calculated from datasets in which residues at one mutated position were fixed. This was done by computing the frequency matrices for the set of antibodies that contained each fixed residue. The full dataset was broken into subsets of sequences that contained the given residue, and these subsets was used to create new PSERMs. Typically, the size of the subsets used to make a fixed residue PSERM were an order of magnitude smaller than the full PSERM datasets.

Fixed residue PSERMs are denoted with a superscript of the fixed amino acid and the fixed position. For example, the PSERM created from sequences with a threonine (T) in the heavy chain Kabat position 55 (H55) was denoted as:

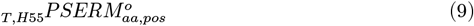

For fixed residue PSERMs, the amino acid and position of the fixed residue was shown as a subscript before PSERM and the superscript *o* represents the output sample used to create the PSERM such as an antigen sort.

Epistatic shifts in the context of a fixed residue were calculated as the Jensen-Shannon divergence of a Boltzmann-like weighting of PSERM amino acid scores with fixed reside PSERM amino acid scores (Starr *et al*. 2022) for a given position. The Boltzmann-like weighting is defined as:

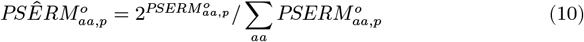

The reweighted PSERMs are denoted with a hat. The Boltzmann-like weighting was done to convert the enrichment ratio profiles into a probability-like distribution to apply the Jansen-Shannon divergence, which quantifies the similarity of two probability distributions. The divergence was calculated between the re-weighted PSERM and the re-weighted fixed residue PSERM for a given position. The divergence was used as a proxy for contextual dependence:

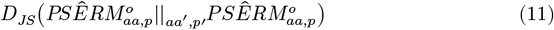

This divergence metric is bounded between zero (identical) and one (dissimilar) and compares how residues differentially enrich at a given mutation site in the context of a fixed residue at a different mutation site. To compute the total contextual dependence of a mutation, the divergence was summed over every other position:

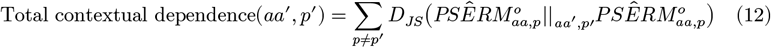

The total contextual dependence is large when a residue causes large epistatic shifts (high *D*_*JS*_ values) at several positions.

### 2.7 Multi-PSERMs

Multi-PSERMs were defined as PSERMs that report the enrichment ratios of multiple residues simultaneously. For example, multi-PSERMs for all possible combinations of both double and triple mutations were created that report the enrichment ratios of each possible pair of sampled residues, denoted as 2-*PSERM*, or triplets of sampled residues, denoted as 3-*PSERM*. Like fixed residue PSERMs, all sequences with a given double or triple set of residues were collected, their frequencies were recorded for both the input and output samples, and their ratios were log transformed to create the multi-PSERMs. This analysis used the same number of sequences as normal PSERMs, but it also distributes the observations into more entries per multi-PSERM due to the combinatorial addition of multiple positions considered. The scoring formula for double and triple sets of residues was analogous to single residue scoring but summed over all pairs or triplets of residues. The triple mutation scoring formula was defined as:

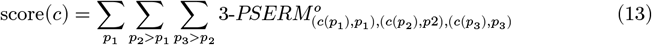

where, *p*_1_, *p*_2_, and *p*_3_ represent the first, second, and third mutated positions of the residue triplet, *c*(*p*_*j*_) represents the amino acid of the scored clone in position *p*_*j*_, and 3-*PSERM*^*o*^ represents the 3-PSERM for output sample *o*.

### 2.8 Statistical analysis

All statistical correlations are Spearman rank correlation coefficients and were computed using SciPy in python (Virtanen *et al*. 2020). Statistical comparison of these Spearman rank coefficients was done by computing the Fisher z-transformed correlation coefficients and performing a Z-test on these transformed coefficients (Raghunathan *et al*. 1996). One-tailed independent t-tests were used to determine statistical significance of differences in mean. For all statistical tests, significance was assumed for tests with a resulting *p*-value < 0.05.

## 3. RESULTS

### 3.1 Clonal PSERM scores are highly reproducible

Toward our goal of developing a new metric for scoring optimal antibody variants in deep sequencing datasets, *in vitro* antibody libraries were sorted for function (i.e., antigen binding) and related properties (e.g., non-specific binding), and the input and output libraries were deep sequenced (**Fig. 1**). We first analyzed the frequency of each antibody variant in the input and output datasets and then converted these frequencies into individual amino acid enrichment ratios in the form of a PSERM. This PSERM is mathematically equivalent to the difference between PSSMs computed from the output sample relative to the input sample. The PSERM is then used to score the variants observed in each sequencing dataset data, and the scores are used to differentiate between different antibody properties.

**Figure 1.**
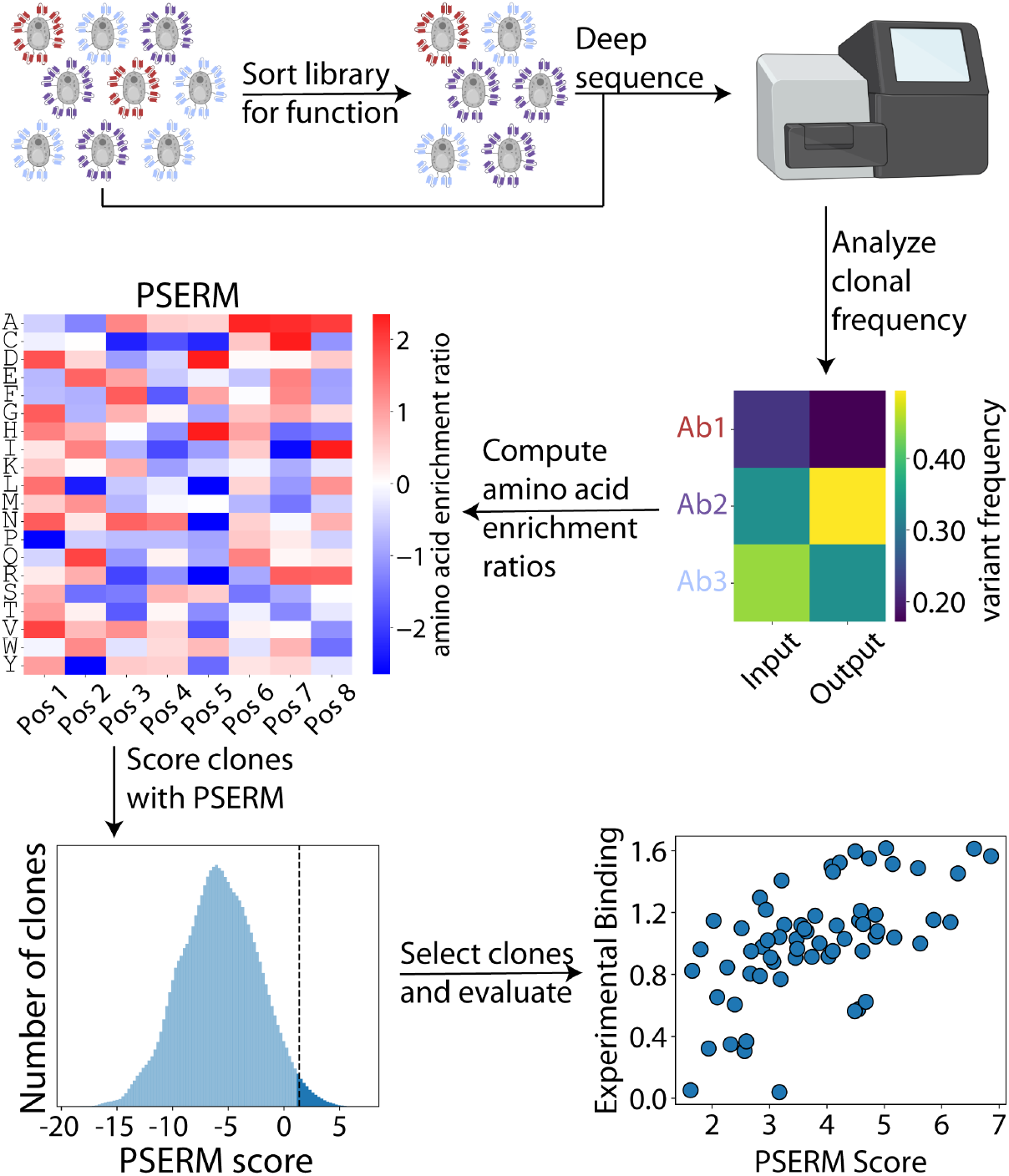
Overview of method for identifying optimal variants in antibody libraries using Position-Specific Enrichment Ratio Matrices (PSERMs). Combinatorial mutagenesis libraries are sorted for desired properties, such as high affinity or low non-specific binding. The input and the output libraries are deep sequenced, and the clonal frequencies are computed. The frequency matrix is transformed into a PSERM, which reports the log_2_ value of the site-specific enrichment ratio of each residue at each mutated site. The PSERM is used to score each observed clone in the library by summing the corresponding site-specific PSERM values for all of the mutated sites. The top scoring clones are selected for analysis and experimental evaluation. Figure created with BioRender.com.

We next sought to compare the reproducibility of sequence scoring metrics – such as frequency, enrichment ratio, and PSERM scores – between replicate sequencing datasets. For example, in **Figure 2A**, the input and output libraries for two replicates of a selection for a lack of antibody self-association are shown for each metric (**Dataset #10**). Notably, the strongest correlation was achieved using PSERM scores (Spearman’s ρ of 0.999), while intermediate correlations were observed for both enrichment ratio (ρ of 0.689) and output frequency (ρ of 0.724). It is interesting that PSERM and enrichment ratio reproducibility is strong considering the low reproducibility of the input frequencies (ρ of 0.242) because both these metrics depend on this data.

**Figure 2.**
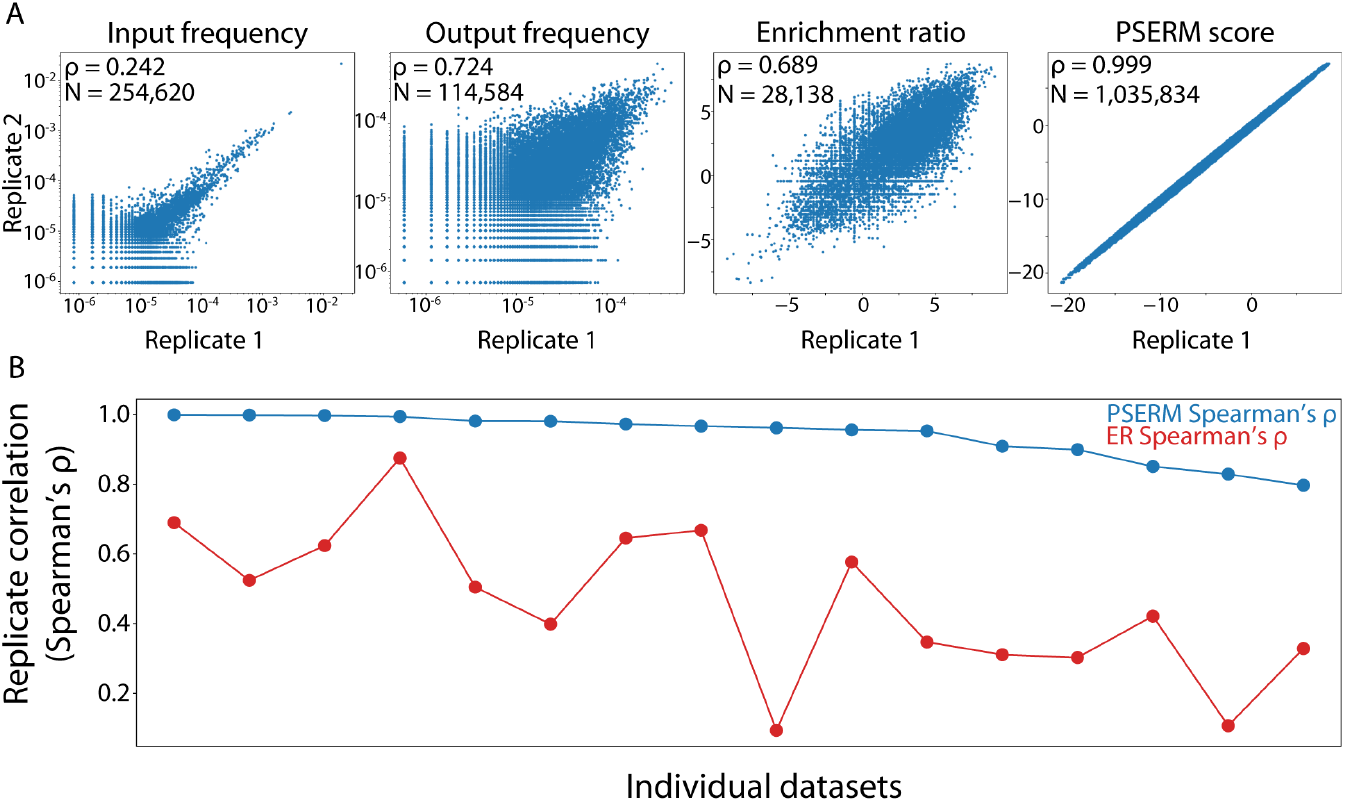
PSERM scores have high reproducibility. (A) Two replicates of deep sequencing data were analyzed separately, and the correlations between the replicates are shown. The first two panels show the frequency of each clone observed in both replicates for input and output samples. The next two panels show the reproducibility of enrichment ratio and PSERM score. (B) Spearman’s ρ correlations are shown for two scoring metrics, namely PSERM and enrichment ratio (ER), for two replicates across 16 different datasets. The datasets are antibody libraries displayed on yeast selected for increased and/or reduced levels of antigen binding, non-specific binding, and self-association. The details of the datasets are provided in **Table S1**, and the details of the sorting process used to generate the library samples is given in **Table S2**.

We next computed these metrics for the other datasets in this study (**Datasets #1-16**) and found that PSERM scores were consistently more correlated between replicates than all other metrics with ρ values ranging from 0.798 to 0.999. On the other hand, the enrichment ratio reproducibility was much more variable with some datasets showing little correlation between replicates (Spearman’s ρ of 0.094), while others showed higher reproducibility (ρ of 0.875) (**Fig. 2B**).

To understand why PSERM scoring is capturing this information more reproducibly, we computed the uncertainty in PSERM score (Eq. 8) and enrichment ratio (Eq. 6). The uncertainty in an enrichment ratio is highest for clones with low observations in the input and output. PSERM scores have a similar uncertainty profile, but the error is based on observations of individual residues at each position rather than observations of a particular antibody sequence. The count of any single residue is at least as much as the least common clone; however, in general, the count of any single mutation is much higher than the count of the least common clone, lowering the uncertainty in any score. As shown in **Figure 3A**, PSERM scores have coefficients of variation that are orders of magnitude lower than the corresponding values for enrichment ratios for the deep sequencing datasets.

**Figure 3.**
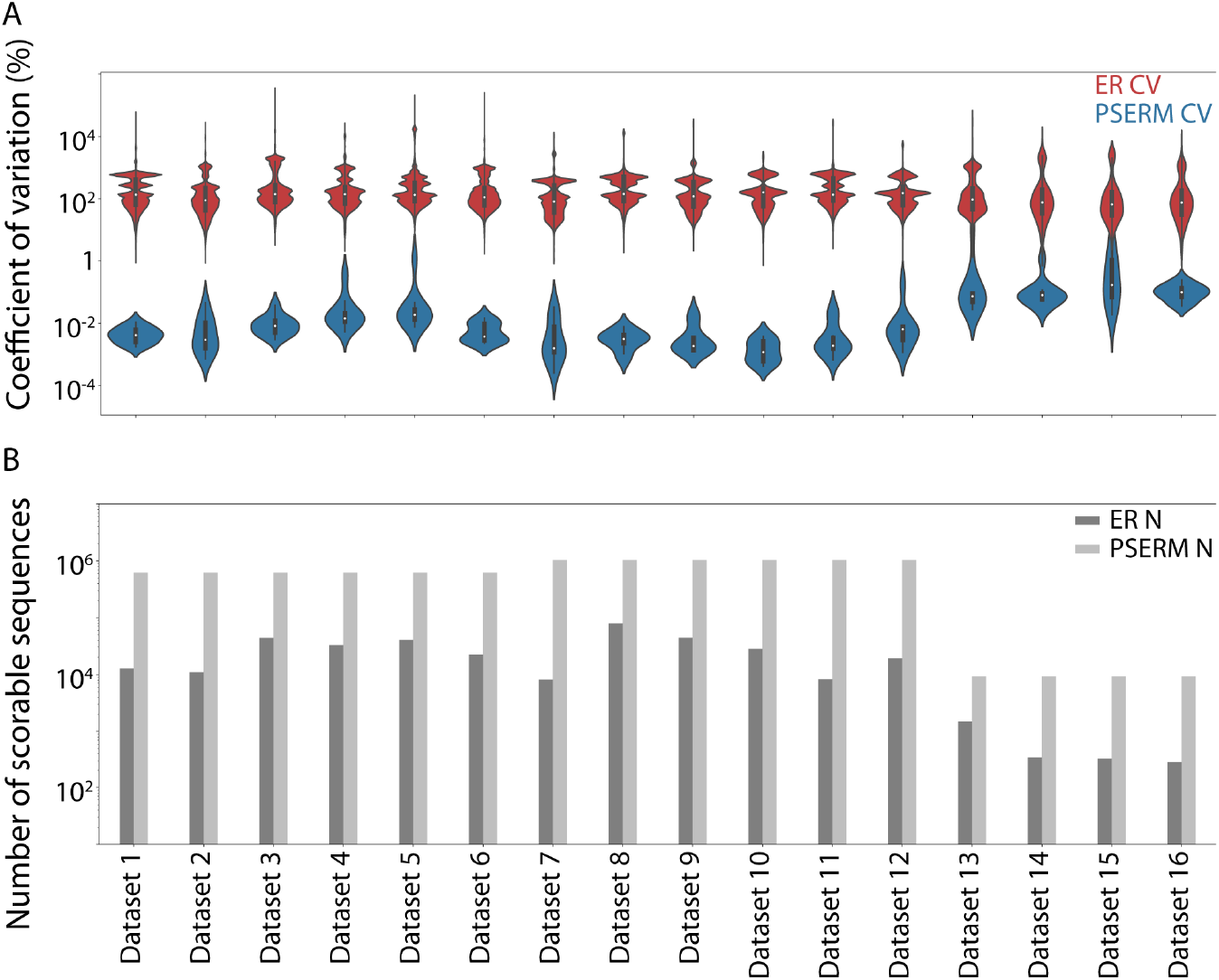
PSERM scoring has lower uncertainty and increased ability to score more clones in libraries relative to conventional methods. (A) The uncertainty in PSERM scores and enrichment ratios (ERs) expressed as the coefficient of variation (CV). The subsets of clones from the datasets that could be assigned an enrichment ratio are shown. (B) The number of scoreable antibody sequences is shown for each dataset for both scoring methods. The details of the datasets are provided in **Table S1**, and the details of the deep sequencing statistics are given in **Table S3**.

We also evaluated the number of antibody variants that could be scored using PSERMs versus enrichment ratios. Computing an enrichment ratio for an antibody variant requires it to be observed in both the input and output datasets, thus limiting the number of scoreable variants, while PSERMs can score any clone given its sequence. Notably, this leads to large increases in the number of scoreable clones (**Fig. 3B**). For example, we observe up to a 125-fold increase in the number of scoreable variants using PSERMs relative to enrichment ratios, which demonstrates the utility of methods that do not require observation of the variants in both the input and output sequencing datasets for the antibody selections. Finally, PSERM scoring uses a non-zero pseudocount (**Eq. 2**) value for calculating the frequency matrices to avoid the potential of taking the logarithm of zero in cases where there are no observations of an amino acid at given site, and we confirmed that PSERM scoring is largely independent on the pseudocount value except in the extreme cases where the number of amino acid observations of a given site is extremely low (e.g., <10 observations; **Fig. S1**).

### 3.2 PSERM scores correlate with experimental binding measurements

We next sought to compare the different metrics with experimental binding measurements. In particular, we compared our computed metrics to experimental datasets reported previously for emibetuzumab mutants (Makowski *et al*. 2022b). These datasets were generated by i) sorting a yeast-display library for both antigen (HGFR) and non-specific (ovalbumin (Makowski *et al*. 2021)) binding, ii) deep sequencing the enriched libraries, and iii) measuring the binding of a set of 125 emibetuzumab variants to antigen and polyspecificity (ovalbumin) reagents (**Datasets #2-3**). We computed the frequency, enrichment ratio, PSSM score, and PSERM score from the deep sequencing data, and compared these metrics to the corresponding normalized binding values for the 125 antibody variants (**Fig. 4**). The antigen and ovalbumin PSERMs are shown in **Figure S2**. Encouragingly, the PSERM scores had the highest correlation coefficient for both reagents (**Fig. 4A-B**). While both PSSM and PSERM yielded high correlation for antigen binding, only PSERM scores had a correlation coefficient >0.5 for ovalbumin binding (**Fig. 4B**).

**Figure 4.**
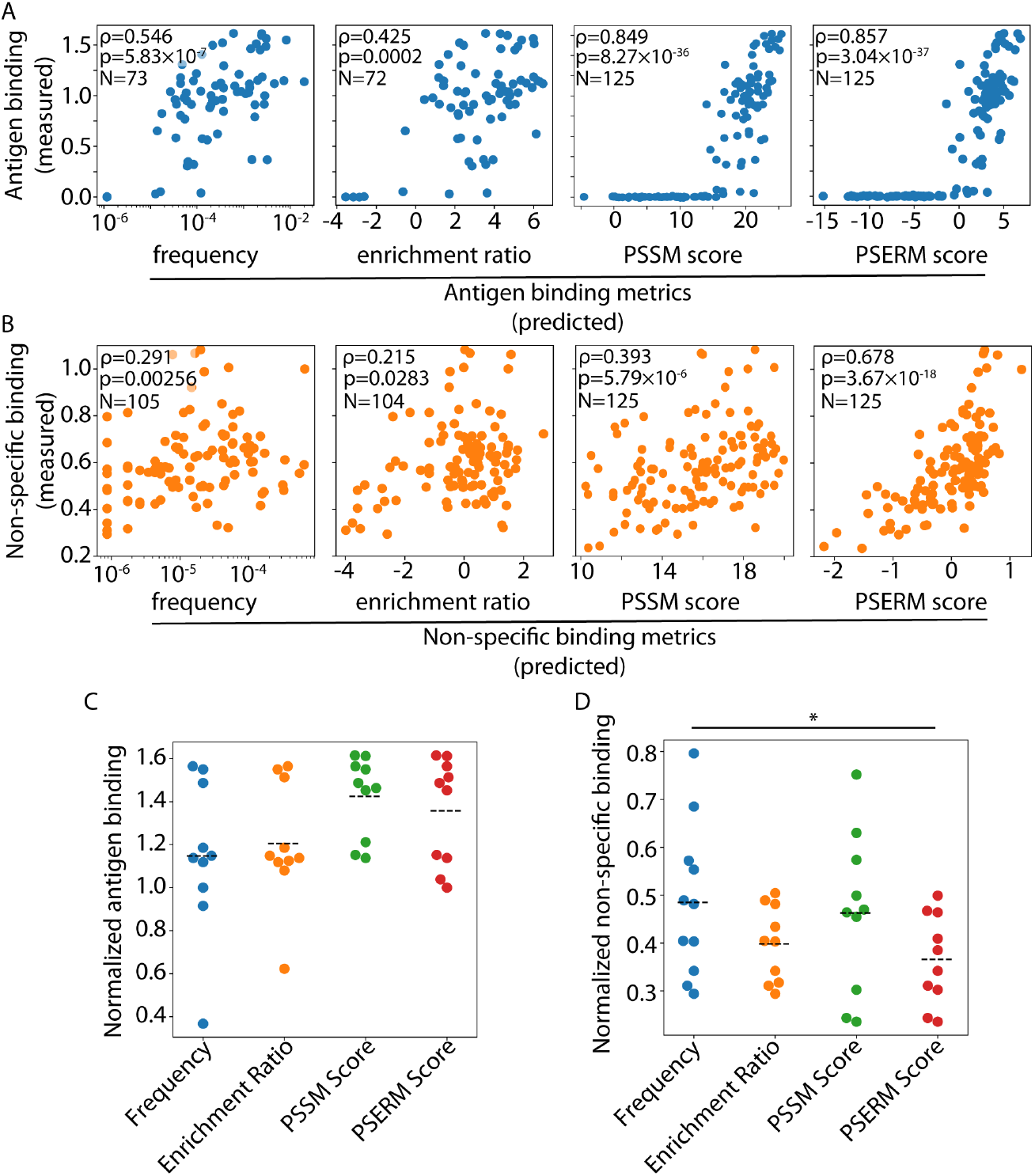
Correlation of predicted binding metrics with experimental measurements. Antibody variants (125 clones) were experimentally evaluated for antigen and non-specific binding and compared to four deep sequencing metrics. (A) Experimentally measured antigen binding was compared to output antigen frequency, enrichment ratio, PSSM score, and PSERM score. (B) Experimentally measured non-specific binding was compared to the same four metrics. (C, D) The experimental binding values of selected clones based on the four different metrics are shown for (C) the 10 highest scoring antigen binding clones and (D) the lowest scoring non-specific binding clones. The black dashed line represents the average binding of the 10 clones. There are 11 clones selected by ovalbumin frequency because they all had the same number of observations. p-values < 0.05 (*).

The PSERM and PSSM scoring methods were able to score all 125 variants, while fewer variants could be scored using frequencies (73 variants for antigen and 105 variants for ovalbumin) and enrichment ratios (72 variants for antigen and 104 variants for ovalbumin). Therefore, we evaluated the correlations between the various metrics and the experimental data using subsets of the 125 clones that could be assigned all four metrics for each binding reagent (**Fig. S3**). This resulted in a set of 72 clones for antigen binding, and 104 clones for ovalbumin binding. While the reduced dataset did lead to a decrease in the correlation coefficients, the PSERM scores were still most strongly correlated for both reagents (Spearman’s ρ >0.6).

We also evaluated the impact of antigen concentration during library enrichment on the PSERM scoring and resulting correlations with experimental antigen-binding measurements (**Fig. S4**). PSERMs were computed for two antigen (HGFR) concentrations, namely 0.1 nM (**Dataset #2**) and 1 nM (**Dataset #1**), and the resulting PSERM scores were compared to the experimental measurements for the 125 antibody (emibetuzumab) variants. The PSERM based on libraries enriched against the lower antigen concentration resulted in higher correlations with the experimental measurements (Spearman’s ρ of 0.857 for 0.1 nM antigen and ρ of 0.825 for 1 nM antigen).

Next, we used the various scoring methods to identify the ten variants that are predicted to have the most favorable binding properties, namely high antigen binding and low non-specific binding, for the emibetuzumab variants (**Fig. 4C**). Notably, the PSERM method gives the best overall performance of selecting variants with high antigen binding and low non-specific binding. For antigen binding, the data rich PSERM and PSSM methods select the highest binding clone and give the highest levels of average antigen binding (1.36x of wild type for PSERM scoring and 1.42x of wild type for PSSM scoring). In contrast, using frequency and enrichment ratio to identify the top ten clones leads to lower average antigen binding (1.15x for frequency and 1.20x for enrichment ratio) and both methods fail to identify the highest binding clone in the dataset. Related analysis of the non-specific binding data reveals similar trends (**Fig. 4D**). Using PSERM scores to identify the ten lowest scoring antibody variants resulted in the lowest average levels of non-specific binding (0.37x of wild type), while the other metrics results in higher average levels of non-specific binding (0.48x for frequency, 0.40x for enrichment ratio, and 0.46x for PSSM scores). Moreover, both PSERM and PSSM scoring selected the low non-specific binding variant.

In many protein selection campaigns, there are often several biophysical properties that need to be optimized simultaneously. Incorporating selections for each property of interest enables the use of multiple PSERMs in concert to select clones with co-optimal properties. Using both the antigen and non-specific binding PSERMs to score the 125 emibetuzumab variants highlights the affinity-specificity tradeoff for this antibody (**Fig. 5**). In general, clones with higher antigen PSERM score also have higher non-specific PSERM scores. Notably, several variants along the affinity-specificity Pareto frontier with positive antigen PSERM scores (**Fig. 5A**) are also located near the experimentally determined Pareto frontier (**Fig. 5B**). The interpretability of the PSERMs reveal the sequence-property relationships that mediate the co-optimal behavior of these antibodies (**Fig. 5C**). For example, the mutation R(H55)G contributes positively to antigen binding and negatively to non-specific binding. In fact, all seven of the Pareto optimal variants have at least two of the co-optimal mutations that improve antigen binding and reduce non-specific binding.

**Figure 5.**
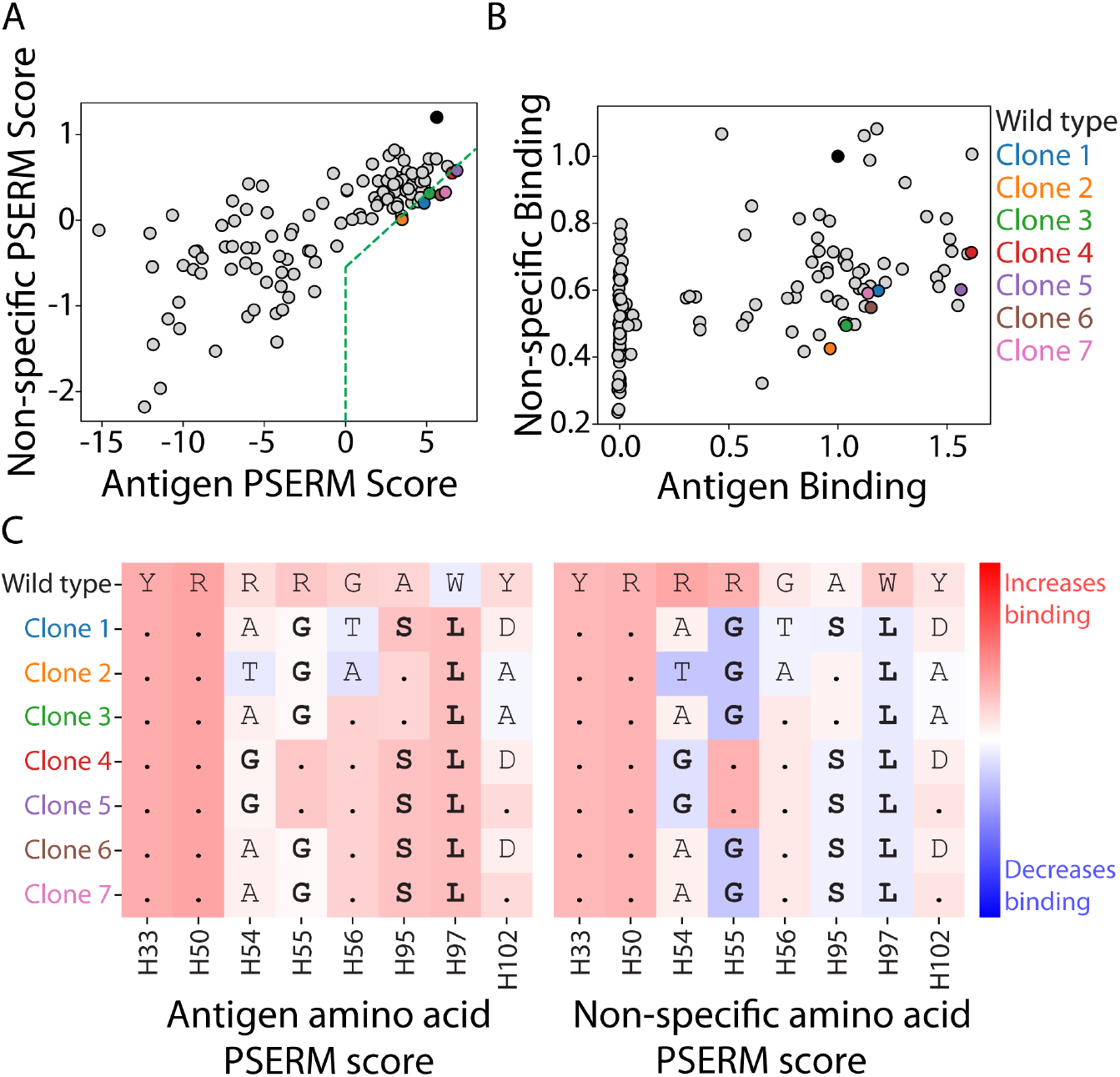
Selection of clones with co-optimal antigen and non-specific binding properties. (A) Antigen and non-specific PSERM scores for the 125 clones that were evaluated experimentally. Seven clones are selected with positive antigen PSERM scores that lie along the Pareto frontier. (B) The clones identified in (A) also lie on, or near, the Pareto frontier identified experimentally. (C) The amino acid PSERM scores of the selected clones. Each row represents a clone, and the entries are colored according to the PSERM value of the mutations. A dot in place of an amino acid represents the wild-type residue in the corresponding sequence. These clones include mutations – shown in bold font – that score positively for antigen binding (red) and negatively for non-specific binding (blue).

We also scored additional emibetuzumab variants that were consistent with the library design but not observed in either in the input or output library samples to evaluate if any of these variants displayed superior antigen or non-specific binding properties (**Fig. S5**). This resulted in scoring an additional 449,853 number of variants, which represents 26.8% of the designed library. Interestingly, we did not find any of the additional clones to have higher predicted antigen binding or lower predicted non-specific binding. Nevertheless, this analysis highlights the utility of PSERM scoring to go beyond what is observed within deep sequencing runs, which is important because library sizes can be larger than the corresponding sequencing depths achieved by common deep sequencing methods.

### 3.3 Mutations with low contextual dependence are scored favorably by PSERMs

We next evaluated potential contextual preferences for certain residues at mutated sites in emibetuzumab using the antigen and non-specific binding datasets (**Datasets #2** and **#3**). Interestingly, we observed very low values for the Jansen-Shannon divergence for the non-specific binding data, suggesting little contextual dependence for non-affinity antibody interactions (**Fig. S6**). However, for antigen binding, we observed variable levels of contextual dependence (**Fig. 6**). For example, the enrichment of different residues at position H102 are unaffected by the presence of tyrosine at position H33 (**Fig. 6A**). This lack of interaction may be due to the positions being in two different CDRs, namely CDRH1 for position H33 and CDRH3 for position H102. However, the enrichment patterns for residues at position H50 strongly differ when position H55 is fixed as threonine (**Fig. 6B**). For example, in this context (threonine at position H55), glutamic acid is strongly enriched at position H50 despite that it is otherwise depleted in the full datasets. This strong dependence may be driven by the proximity of these two positions in the same CDR (CDRH2).

**Figure 6.**
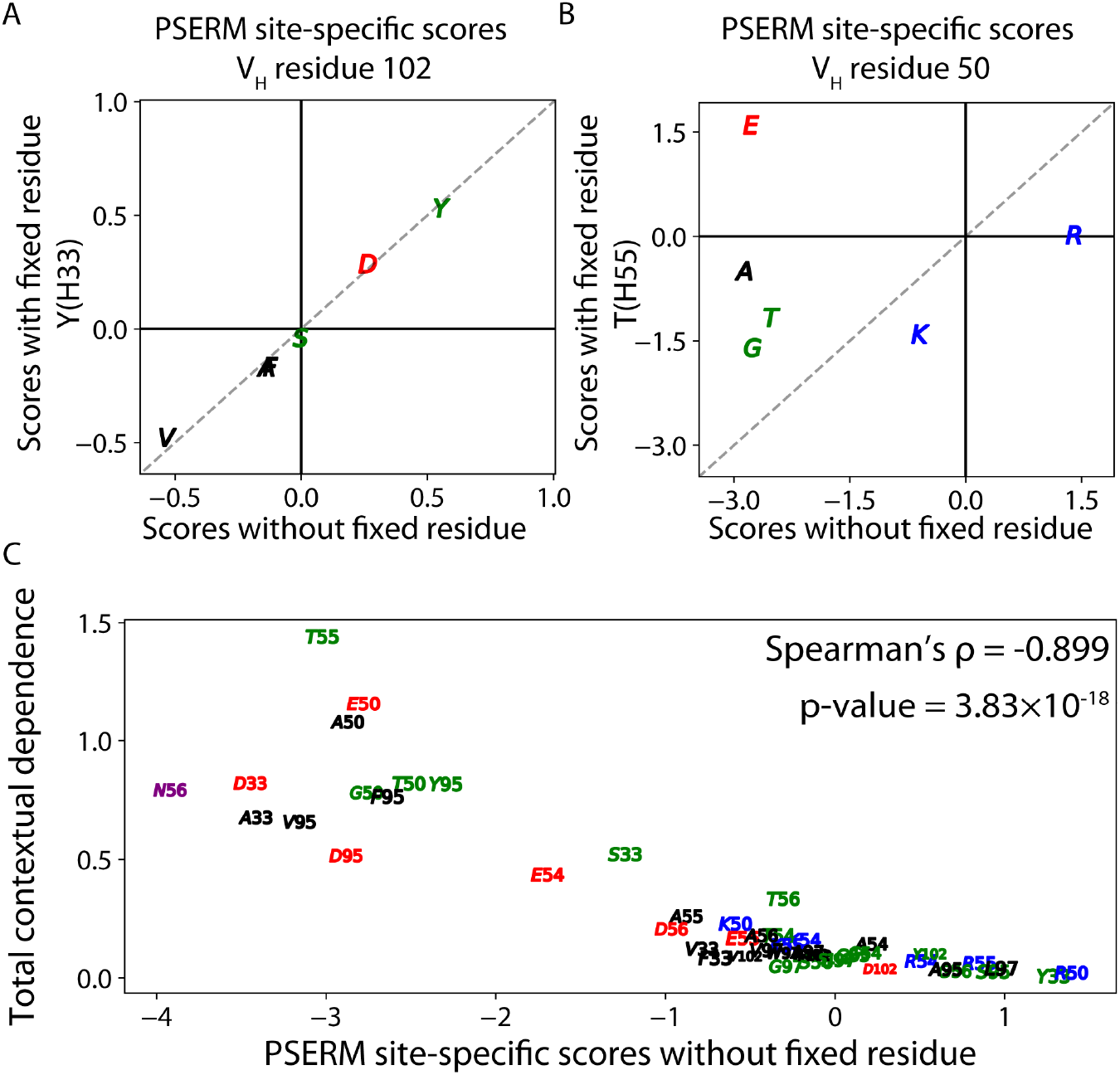
Mutations with low contextual dependence contribute favorably to PSERM scores. (A-B) PSERMs were generated first by fixing a residue at a specific position, namely (A) Y(H33) or (B) T(H55). Next, these fixed residue PSERMs were used to score residues at a second site, namely (A) H102 or (B) H50. Finally, these scores were compared to those generated with PSERMs that do not have a fixed residue to evaluate contextual dependence. This reveals that that the site-specific scores for residues at (A) H102 are weakly impacted by the fixed [Y(H33)], while scores for (B) H50 are strongly impacted by the fixed residue [T(H55)]. (C) Total contextual dependence of residues at the mutated sites negatively correlates with PSERM site-specific scores. The total contextual dependence is defined as the sum of the Jansen-Shannon divergence of the mutation with all other positions, as defined in **Eq. 12**.

To further evaluate the contextual dependence of each residue observed at each mutated site, we compared the PSERM site-specific scores without fixed residues to their corresponding total contextual dependence values, the latter of which is defined as the Jansen-Shannon divergence of each residue summed over all other non-fixed positions (**Fig. 6C**). For example, we fixed the mutation R(H55)T and summed the Jansen-Shannon divergence values for all other residues at all mutated positions, which resulted in the highest observed positive value. For the same mutation, we observed a negative site-specific PSERM score. This behavior was observed more broadly for the other mutations, as those mutations with negative (unfavorable) site-specific PSERM scores generally displayed high contextual dependence values and mutations with positive (favorable) site-specific scores generally displayed low contextual dependence.

Our observations that certain amino acids preferred to be paired together, such as residues T(H55) and E(H50), led us to test if PSERM scores would be more strongly correlated with the corresponding experimental data if they reflected the enrichment of combinations of residues rather than each individual residue on its own. To do this, we constructed multi-PSERMs that evaluated every pair or triplet of amino acids in each non-redundant combination (**Fig. S7**). Scoring clones with these multi-PSERMs should allow for second or third order non-linear effects to be reflected in the overall score. Notably, increasing the number of residues scored together had little impact on the correlation for either antigen or non-specific binding (**Fig. 7**). We only observed a slight increase in the correlation constant for non-specific binding (0.0063 increase to Spearman’s ρ for 3-residue scoring), although this was not statistically significant.

**Figure 7.**
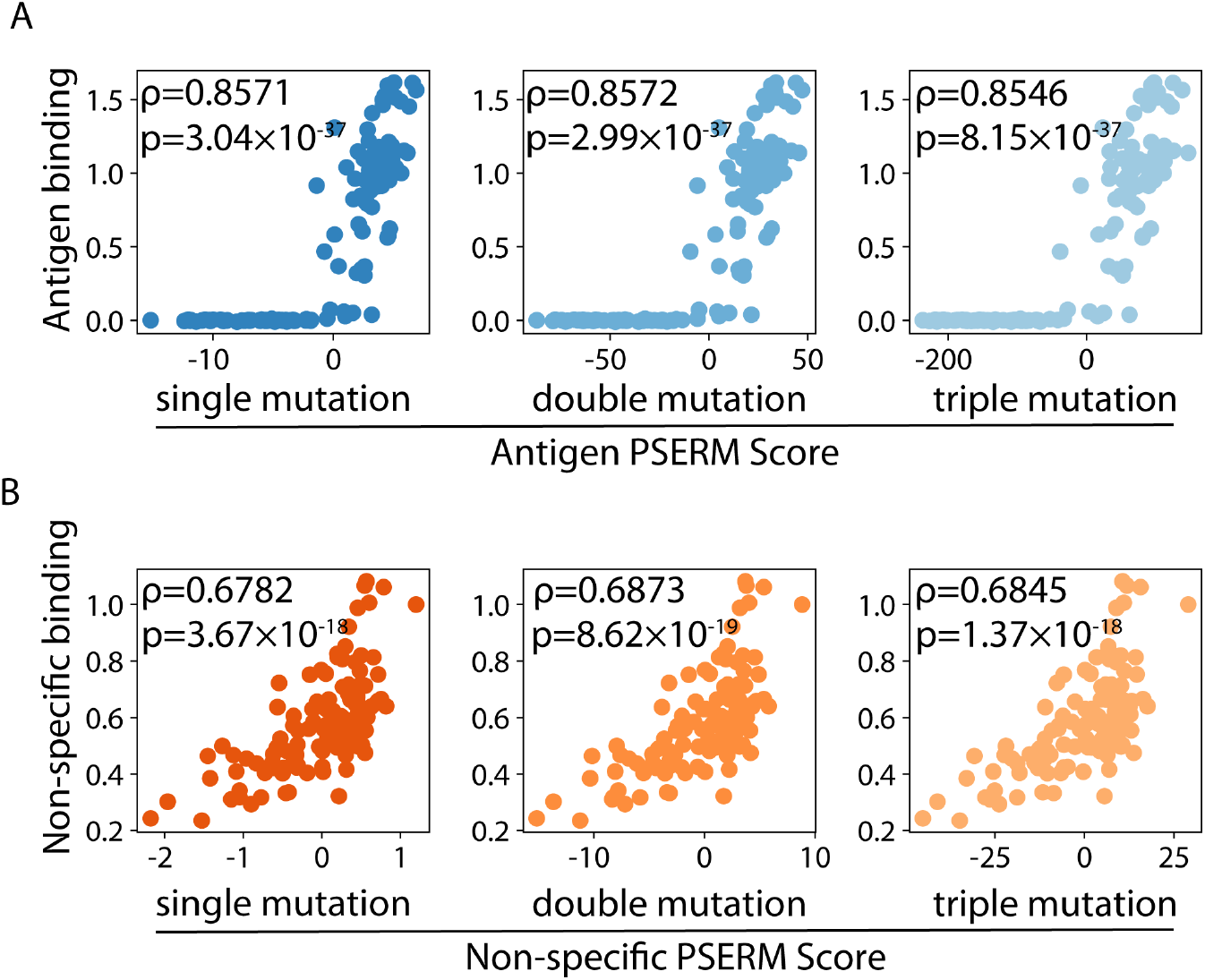
PSERM scoring can be adapted to include contextual dependence. (A) Experimentally measured antigen binding is compared to PSERM scores that (left) do not consider contextual dependence as well as those that consider (middle) co-enrichment of pairs of residues or (right) co-enrichment of triplets of residues. (B) The same three scoring methods and corresponding experimental datasets are shown for non-specific binding. For (A), the multi-PSERMs that consider enrichment of pairs and triplets of residues are further explained in **Figure S7**.

## 4. DISCUSSION

Traditionally, after a selection campaign, candidates for further evaluation are chosen from Sanger sequencing results of the output variants (Rouet *et al*. 2018). This low throughput method allows for testing of tens to hundreds of clones. However, with the advent of deep sequencing, it is now routine to generate sequences of millions of variants, including rare clones that only are observed extremely infrequently, such as once in a million reads. Most deep sequencing campaigns have utilized simple metrics such as frequency or enrichment ratio to select clones for further analysis, but these methods fail to use the vast majority of the sequencing data and have large uncertainties that can lead to limited ability to identify optimal clones. In this work, we demonstrate that the PSERM score de-couples the score of a clone from its number of observations, allowing for even the rarest clone to be scored with much less uncertainty than traditional scoring methods.

Because the abundance or enrichment ratio of a variant is not always indicative of the selected antibody or protein property of the variant, methods that de-couple a variant score from such metrics are of great interest (Reich *et al*. 2015; Adams *et al*. 2016; Kelil *et al*. 2021; Saka *et al*. 2021). We have shown that our simple method of PSERM scoring outperforms traditional frequency and enrichment-based scoring of antibody variants from deep sequencing data, in both the average binding improvement and correlation with affinity and non-specific binding properties. The performance of the PSERM method is notable given that only deep sequencing data is used from single input and output library samples without the need for multiple antigen or polyreactivity reagent concentrations and/or multiple gates during the sorting process (Reich *et al*. 2015; Adams *et al*. 2016; Jenson *et al*. 2018). Further, because this method utilizes all the data in the deep sequencing datasets, the PSERM scores have extremely low uncertainty and show high reproducibility between sequencing replicates.

The improved performance of PSERM scoring relative to PSSM scoring deserves further consideration. PSSM scoring is a much more comprehensive method than only scoring protein variants based on their output frequency, yet the two approaches are similar in that they only utilize information observed in the output samples after selection. This inherently makes them dependent on the frequencies of protein variants in the input library, and this information is not considered in either scoring method. Conversely, PSERM scoring is a much more comprehensive method than enrichment ratio scoring of a given observed protein variant, yet the two approaches are also similar in that they use information from both the input and output samples. It is likely that the superior performance of PSERMs relative to PSSMs is simply due to the increased information content (which comes from the input samples) that is also encoded into the PSERMs.

It is important to note that the antibody sequence datasets presented in this work are from antibody maturation campaigns with mutations only in the CDRs and represent a focused sampling of sequence space. The libraries in this work also lack V_H_ or V_L_ framework, V_H_/V_L_ pairing, and CDR length diversity, and the mutations in each library are not expected to change the epitope. We expect that PSERM scoring will also be useful for naïve antibody libraries that have much greater diversity, but this may require developing PSERMs that are specific for each framework, V_H_/V_L_ pairing and CDR length. The success of this approach will likely be dependent on sequencing depth given the more fragmented nature of this analysis.

Another benefit of PSERM scoring is the ease of using this approach to understand sequence-function relationships for each selected property, such as antibody affinity and non-specific binding. It should be noted that many studies have used Deep Mutational Scanning (DMS) to explore such sequence-function relationships for different protein properties (Fowler *et al*. 2014; Bloom 2015; Abriata *et al*. 2016; Rubin *et al*. 2017), including for enzyme solubility and fitness (Klesmith *et al*. 2017; Wrenbeck *et al*. 2019), protein-protein binding, and antibody-antigen binding (Forsyth *et al*. 2013; Koenig *et al*. 2015; Woldring *et al*. 2015; Warszawski *et al*. 2019). However, DMS typically does not involve multi-site mutations, while each of the datasets in this study only involved multi-site mutation libraries. Moreover, DMS cannot be directly used to score multi-site mutant variants without developing corresponding models, while our PSERM approach does not require any model development.

One underlying assumption of PSERM scoring is that each residue acts independently and thus the enrichment ratios of each residue can be summed across multiple positions to score multi-site mutant variants. In fact, the combination of enrichment ratios for individual residues in multi-site mutant libraries has been previously reported (Fowler *et al*. 2010; Reich *et al*. 2015; Naftaly *et al*. 2018). For example, one study used single-site libraries and phage display to define sequence-function relationships for a small protein (WW) domain and found that the product of two single-mutation enrichment ratios was strongly correlated with the enrichment ratios of the corresponding double mutants (Fowler *et al*. 2010). A second study used multi-site mutant libraries of a BH3 peptide, along with yeast-surface display and multi-gate FACS sorting, to identify optimal peptide variants (Reich, Dutta and Keating 2015). One of their scoring metrics was the product of enrichment ratios at multiple sites, which is closely related to our PSERM scores, but they only used this metric to rank variants and failed to demonstrate whether it is directly correlated with peptide binding properties. Our work builds on these important studies for peptides and small proteins and demonstrates the power of PSERMs for accurately predicting optimal multi-site antibody variants without the need for complex sorting methods or models.

Because PSERM scoring removes the observational requirements of a protein variant for scoring, any sequence in the designed library could be scored even if it was never observed (**Fig. S5**). A simple application of this idea would be to design and test a protein sequence composed of the highest scoring amino acid at each position. Further analysis with PSERMs describing multiple protein properties could also be performed to select co-optimal variants. In fact, similar work has achieved this with unrelated models to design highly specific peptides to different members of the B cell lymphoma 2 family (Case *et al*.; Jenson *et al*. 2018). This type of optimization could easily be performed using PSERMs to identify optimal variants, even if unseen in the sequencing data.

Epistatic effects are widely known to occur and can lead to non-additive effects (McLaughlin *et al*. 2012; Otwinowski *et al*. 2018; Starr *et al*. 2022). For example, one report evaluated methods for scoring protein-peptide affinities for three members of the Bcl-2 family, and found that non-linear models were able to better predict the affinities than corresponding linear models (Jenson *et al*. 2018). Even though we observed preferential enrichment of pairs of amino acids within the datasets explored in this work, multi-residue scoring did not lead to significant improvements over single residue scoring. While there are many potential reasons for such behavior, one plausible explanation is that the enrichment ratios for each residue at each single site already reflect at least some of the non-additive effects because they are computed from many observations that capture such contextual dependence. This speculative idea will need to be further tested in the future.

Co-optimization is an important and common issue within protein engineering campaigns (Klesmith *et al*. 2017; Rabia *et al*. 2018; Stimple *et al*. 2020). Antibodies routinely have trade-offs between antigen affinity and other properties, such as stability and/or non-specific binding (Julian *et al*. 2017; Shehata *et al*. 2019; Makowski *et al*. 2022b). The need for a simple metric to score variants for multiple properties simultaneously is critical for identifying clones with Pareto (co-optimal) behavior. Our scoring method is well suited for this task, as we demonstrated scoring of antibody variants for both antigen and non-specific binding and showed that these scores were predictive of co-optimal variants along the experimental Pareto frontier. This method could be expanded to select antibodies with various co-optimal sets of properties, including broadly neutralizing variants against multiple viruses (Sok *et al*. 2018; Magar *et al*. 2021) or species cross-reactive antibodies against orthologs of the same target for clinical translation (Ørstrup *et al*. 2019; Li *et al*. 2020).

In summary, we have demonstrated a robust and simple method for scoring antibody variants in deep sequencing datasets using the Position-Specific Enrichment Ratio Matrix (PSERM). This method uses information from all protein variants sequenced in the input and output datasets after selection to achieve a highly reproducible and accurate metric for predicting experimental binding measurements. This method gives a deeper understanding of which residues contribute to improved functional and biophysical properties and can be used for co-optimal variant selection. Further, the scoring algorithm can be easily modified to include multi-residue scoring in datasets with high levels of epistasis or non-additive effects. While this analysis has been presented as an antibody engineering project, this methodology should be broadly applicable to diverse protein engineering studies.

## 5. ACKNOWLEDGEMENTS

We would like to thank the Advanced Genomics Core from the University of Michigan Biomedical Research Core Facilities for sequencing the datasets in this work and for their help troubleshooting the experiments.

## 6. FUNDING

This work was supported by the National Institutes of Health (RF1AG059723 and R35GM136300 to P.M.T.) and National Science Foundation (CBET 1159943, 1605266 and 1813963 to P.M.T., Graduate Research Fellowship to M.D.S.), the Albert M. Mattocks Chair (to P.M.T).

## 7. DATA AVAILABILITY

All deep sequencing datasets as well as the jupyter notebooks containing the analyses to generate the figures can be found at the github repository: https://github.com/Tessier-Lab-UMich/PSERM_paper.

## Notes

https://github.com/Tessier-Lab-UMich/PSERM_paper

### Competing Interest Statement

The authors have declared no competing interest.

