## Supplemental information for "Position-Specific Enrichment Ratio Matrix scores predict antibody variant properties from deep sequencing data"

### 1. Supplemental Tables

**Table S1. Summary of antibody library deep sequencing datasets and replicate statistics.** The three projects are for three different antibody libraries for antibodies specific for hepatocyte growth factor receptor (**Project 1**), platelet derived growth factor BB (**Project 2**) and Aβ fibrils (**Project 3**). The sample codes for each library indicate the type of selection that was performed, including Ag1 for antigen at 1 nM, Ag01 for antigen at 0.1 nM, OvaP for positive ovalbumin selection, OvaN for negative ovalbumin selection, PSR_P for positive polyspecificity reagent (PSR) selection, PSR_N for negative PSR selection, AgP for positive antigen selection, AgN for negative antigen selection, QDP for positive selection against lenzilumab conjugated to quantum dots, QDN for a negative selection against lenzilumab conjugated to quantum dots, EP for positive enzyme selection, EN for negative enzyme selection, R3 for round 3 output, R5 for round 5 output, R6 for Round 6 output and R7 for round 7 output. The Spearman's ρ values are given for the correlations between two deep sequencing replicates for two scoring metrics, namely PSERM and enrichment ratio (ER), which are reported in graphical form in **Figure 2**. Finally, the number of scorable sequences by each metric (PSERM and ER) are also given, which is also reported in graphical form in **Figure 3**.

| Project | Dataset | Sample | PSERM Spearman's ρ | ER Spearman's ρ | PSERM N | ER N |
|---|---|---|---|---|---|---|
| 1 | Dataset 1 | Ag1 | 0.899 | 0.30 | 615,575 | 12,820 |
| 1 | Dataset 2 | Ag01 | 0.953 | 0.35 | 615,575 | 10,951 |
| 1 | Dataset 3 | OvaP | 0.962 | 0.09 | 615,575 | 44,175 |
| 1 | Dataset 4 | OvaN | 0.909 | 0.31 | 615,575 | 33,091 |
| 1 | Dataset 5 | PSR_P | 0.829 | 0.11 | 615,575 | 40,823 |
| 1 | Dataset 6 | PSR_N | 0.797 | 0.33 | 615,575 | 22,438 |
| 2 | Dataset 7 | AgP | 0.994 | 0.87 | 1,035,834 | 8,116 |
| 2 | Dataset 8 | AgN | 0.980 | 0.40 | 1,035,834 | 79,436 |
| 2 | Dataset 9 | QDP | 0.998 | 0.52 | 1,035,834 | 44,468 |
| 2 | Dataset 10 | QDN | 0.999 | 0.69 | 1,035,834 | 28,138 |
| 2 | Dataset 11 | EP | 0.997 | 0.62 | 1,035,834 | 8,271 |
| 2 | Dataset 12 | EN | 0.981 | 0.50 | 1,035,834 | 19,307 |
| 3 | Dataset 13 | R3 | 0.851 | 0.42 | 9,366 | 1,504 |
| 3 | Dataset 14 | R5 | 0.956 | 0.58 | 9,366 | 348 |
| 3 | Dataset 15 | R6 | 0.973 | 0.65 | 9,366 | 334 |
| 3 | Dataset 16 | R7 | 0.966 | 0.67 | 9,366 | 286 |

**Table S2. Summary of sorting conditions and reagents.** For each project and library sample, the relevant sorting methods, reagents, incubation times, temperatures, and buffer conditions are supplied. The details regarding the library sample names are given in **Table S1**.

| Project | Sorting method | Dataset | Sample | Reagent | Incubation time | Incubation temperature | Buffer conditions |
|---|---|---|---|---|---|---|---|
| 1 | FACS | Dataset 1<br>Dataset 2 | Ag1<br>Ag01 | 1 nM HGFR-Fc<br>0.1 nM HGFR-Fc | 3 h | RT | 1x PBS + 0.1% BSA + 1% milk |
| | | Dataset 3<br>Dataset 4 | OvaP<br>OvaN | 260 μg/mL Ovalbumin | 3 h | RT | 1x PBS + 0.1% BSA |
| | | Dataset 5<br>Dataset 6 | PSR_P<br>PSR_N | 130 μg/mL soluble membrane proteins | 20 min | Ice | 1x PBS + 0.1% BSA |
| 2 | FACS | Dataset 7<br>Dataset 8 | AgP<br>AgN | 5 nM PDGF-BB | 3 h | RT | 1x PBS + 0.1% BSA + 1% milk |
| | | Dataset 9<br>Dataset 10 | QDP<br>QDN | 100x dilution of QD-lenzilumab | 1 h | RT | 0.1x PBS + 0.1% BSA |
| | | Dataset 11<br>Dataset 12 | EP<br>EN | 500 nM LPLA2 | 1.5 h | RT | 0.1x PBS + 0.1% BSA |
| 3 | MACS | Dataset 13<br>Dataset 14<br>Dataset 15<br>Dataset 16 | R3<br>R5<br>R6<br>R7 | $10^7$ Aβ coated dynabeads | 3 h | RT | 1x PBS + 0.1% BSA + 1% milk |

**Table S3. Summary of deep sequencing reads and replicate statistics.** For each project and dataset, the number of total reads and unique sequences observed for each replicate are given. A description of sample names is given in **Table S1**.

| Project | Theoretical library size | Antibody format | Dataset | Sample | Number of reads replicate 1 | Number of unique sequences replicate 1 | Number of reads replicate 2 | Number of unique sequences replicate 2 |
|---|---|---|---|---|---|---|---|---|
| 1 | 1,679,616 | single-chain Fab | NA | Input | 1,124,801 | 534,622 | 422,021 | 275,256 |
| | | | Dataset 1 | Ag1 | 852,614 | 68,642 | 362,501 | 49,131 |
| | | | Dataset 2 | Ag01 | 479,012 | 45,135 | 407,575 | 56,602 |
| | | | Dataset 3 | OvaP | 784,516 | 375,435 | 377,648 | 218,102 |
| | | | Dataset 4 | OvaN | 1,003,960 | 339,053 | 329,373 | 193,742 |
| | | | Dataset 5 | PSR_P | 1,223,220 | 527,141 | 434,121 | 184,603 |
| | | | Dataset 6 | PSR_N | 332,532 | 165,968 | 349,308 | 193,221 |
| 2 | 1,555,200 | single-chain Fab | NA | Input | 1,517,232 | 842,264 | 1,289,726 | 806,200 |
| | | | Dataset 7 | AgP | 2,018,188 | 192,535 | 1,744,902 | 203,482 |
| | | | Dataset 8 | AgN | 2,154,312 | 1,002,697 | 1,386,191 | 733,310 |
| | | | Dataset 9 | QDP | 1,980,461 | 813,486 | 1,473,517 | 686,594 |
| | | | Dataset 10 | QDN | 2,093,513 | 447,737 | 1,640,062 | 361,305 |
| | | | Dataset 11 | EP | 608,639 | 195,398 | 488,645 | 139,865 |
| | | | Dataset 12 | EN | 577,740 | 307,720 | 499,735 | 271,566 |
| 3 | 10,240,000,000,000 | single-chain Fv | NA | Input | 808,243 | 17,643 | 1,066,991 | 16,760 |
| | | | Dataset 13 | R3 | 883,327 | 5,920 | 1,237,003 | 6,885 |
| | | | Dataset 14 | R5 | 965,836 | 1,697 | 1,014,754 | 2,084 |
| | | | Dataset 15 | R6 | 936,163 | 1,497 | 1,124,647 | 1,762 |
| | | | Dataset 16 | R7 | 968,311 | 1,590 | 992,698 | 1,672 |

### 2. Supplemental Figures

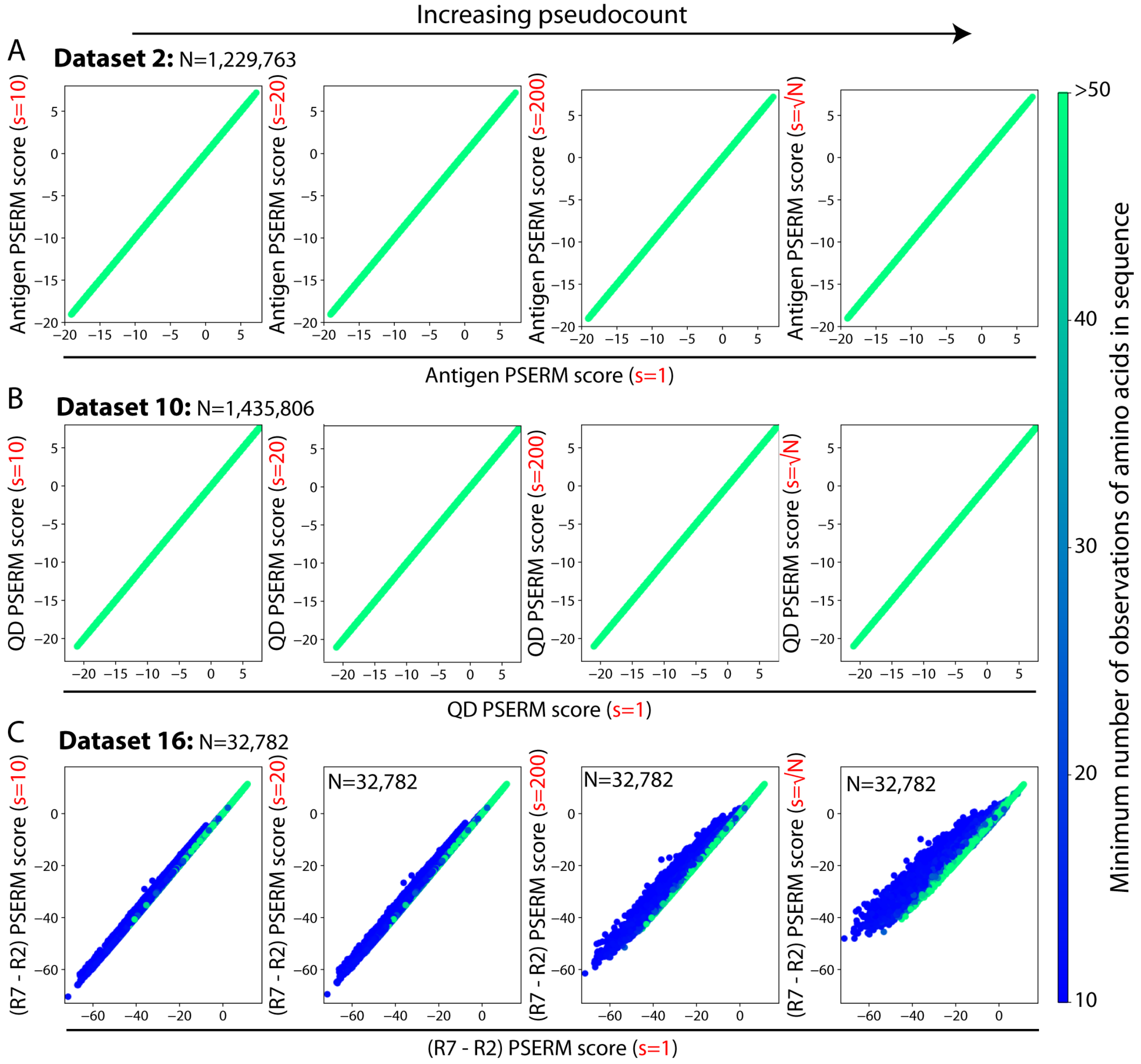


**Figure S1. Effect of pseudocount value on PSERM scoring.** The PSERM score of each observed clone is compared for different pseudocount values across three antibody engineering projects, namely (A) **Dataset #2**, (B) **Dataset #10**, and (C) **Dataset #16**. The x-axis in each plot is the original PSERM score computed with pseudocount (s) of 1. The y-axis of each graph represents the score of the clones with increasing pseudocount values (s = 10, 20, 200, √N). The color of the data point indicates the number of observations of the minimally observed amino acid at any mutated position within a given sequence.

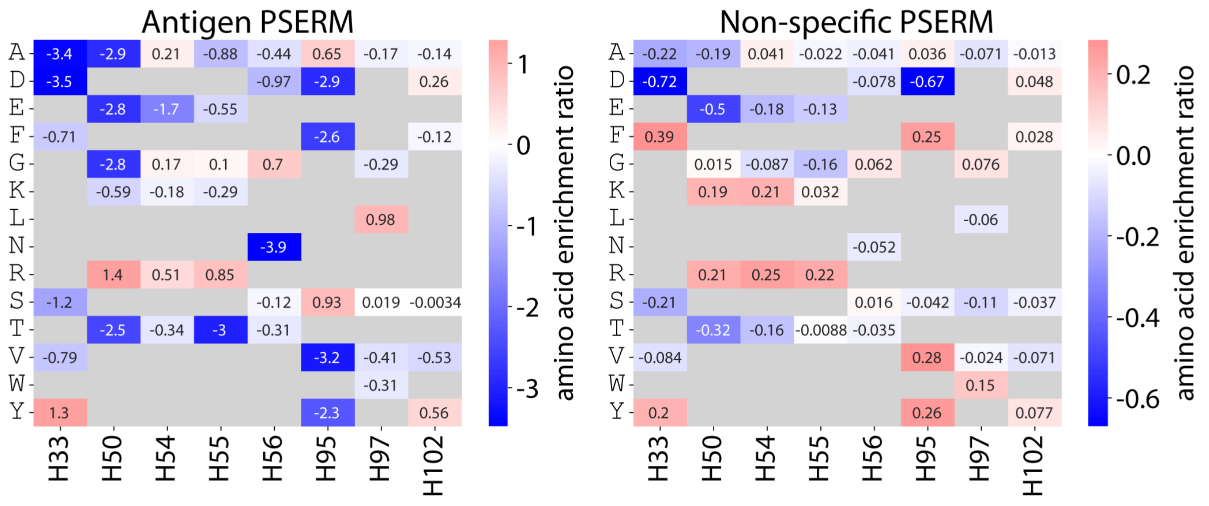


**Figure S2. Antigen and non-specific PSERMs.** PSERMs and the corresponding site-specific enrichment ratios of each sampled residue for (A) antigen (**Dataset #2**) and (B) non-specific binding (**Dataset #3**) selections. Amino acids that are colored red are enriched and correlate with improved binding, while those that are colored blue are depleted after sorting and correlated with decreased binding.

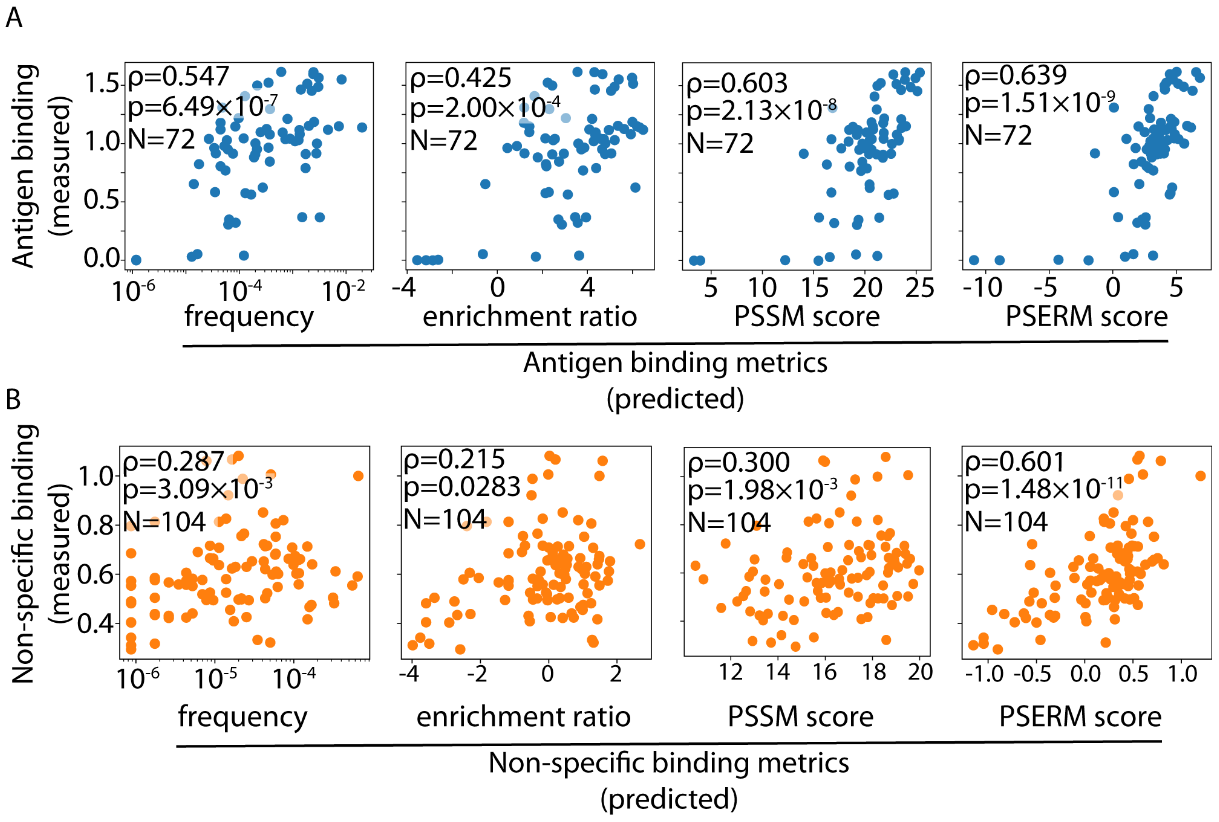


**Figure S3. Correlation of predicted binding metrics with experimental measurements for a subset of the antibody variants.** Correlations between experimental data for a subset of the 125 antibody variants, namely (A) 72 for antigen binding and (B) 104 for non-specific binding, and the corresponding scoring metrics. This figure is based on Figure 4 but only contains a subset of variants that could be scored by all four metrics.

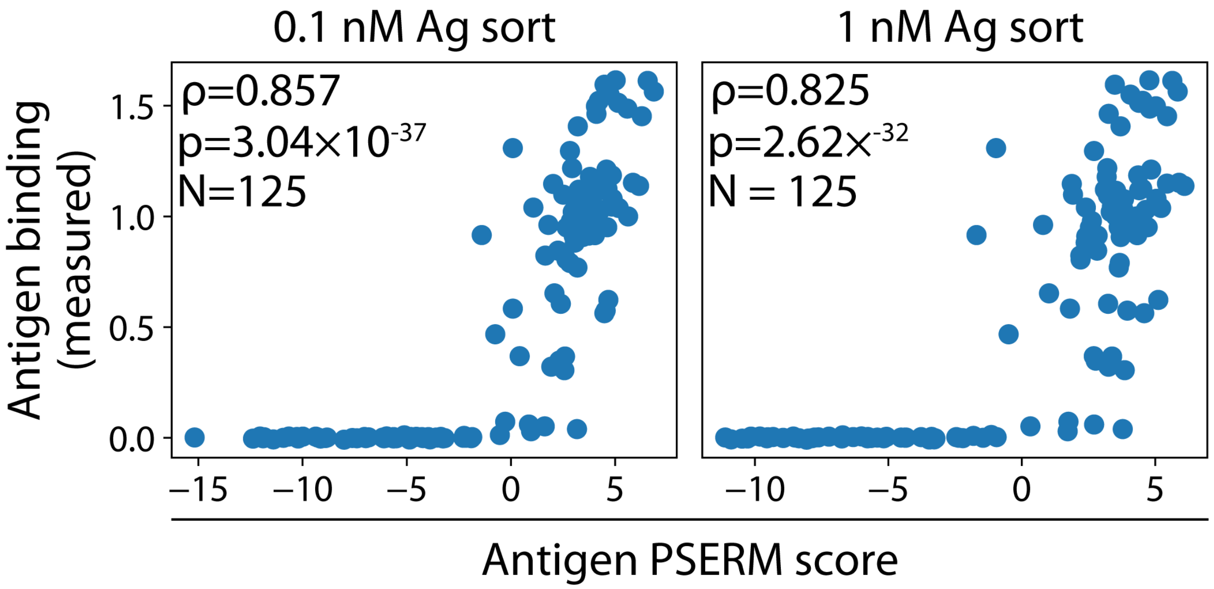


**Figure S4. Lower selection pressure reduces PSERM correlation with experimental measurements of antigen binding.** PSERMs were computed from antigen selections performed with two different antigen concentrations, namely (A) 0.1 nM HGFR (**Dataset #2**) and (B) 1 nM HGFR (**Dataset #1**), and the PSERM scores were compared to experimental measurements of antigen binding.

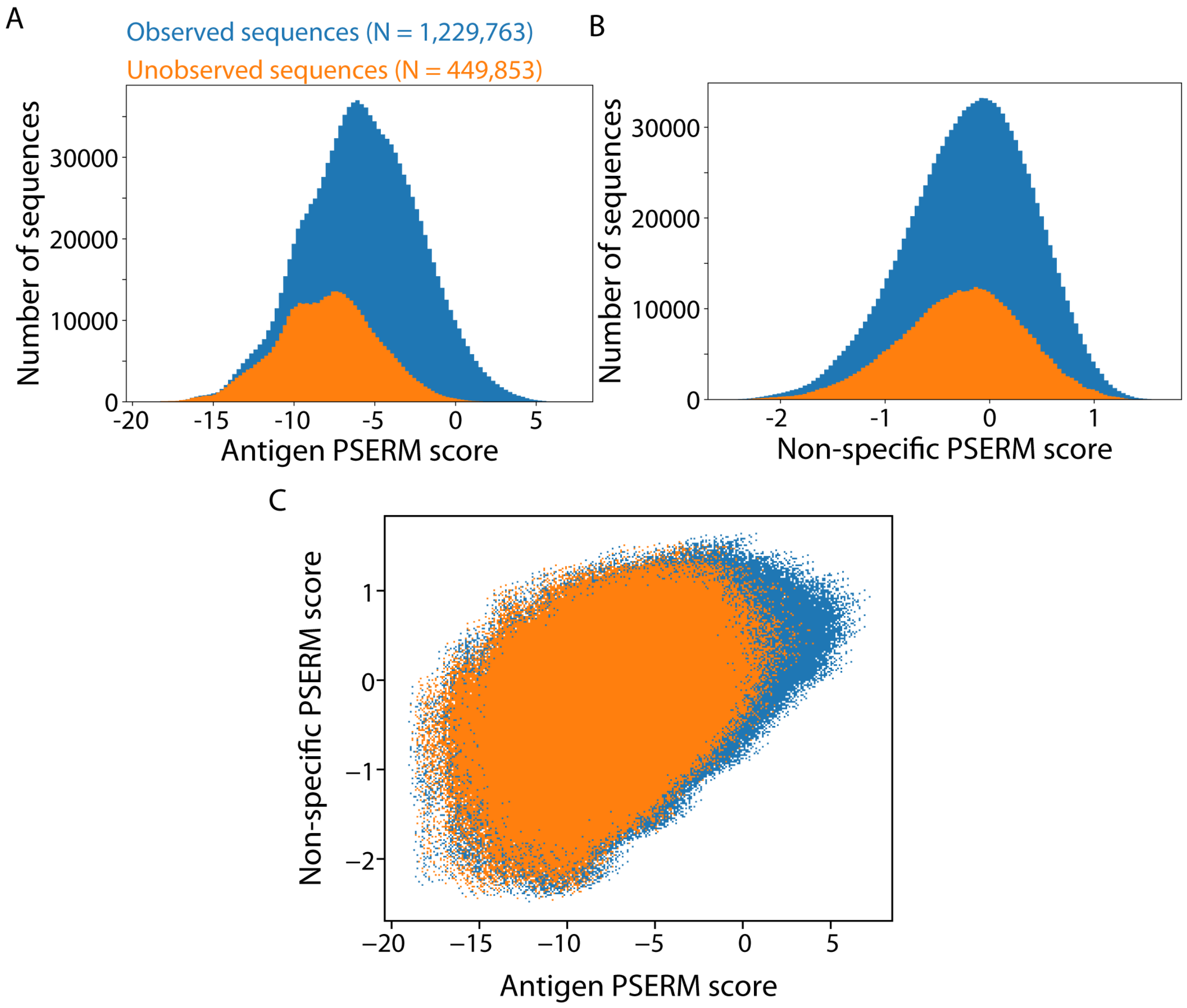


**Figure S5. PSERM scoring of unobserved clones.** PSERM score distributions of observed and unobserved clones for (A) antigen binding (**Dataset #2**) and (B) non-specific (**Dataset #3**) binding. (C) Pareto plot showing the trade-off between antigen and non-specific binding of all clones designed in the library.

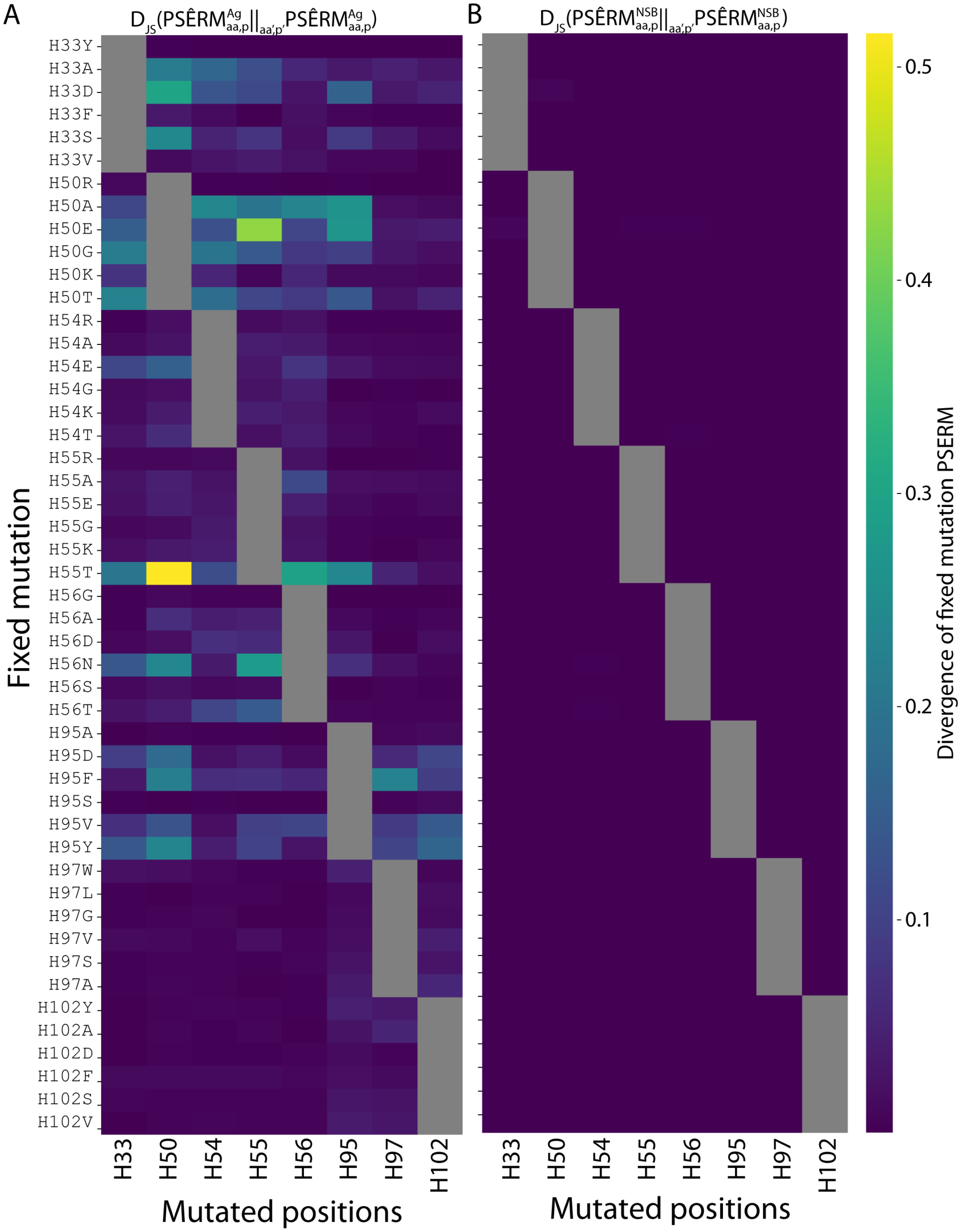


**Figure S6. Contextual dependence of each mutation.** For each mutation, divergence values were calculated, as defined in **Equation 11**, to evaluate if fixing one residue at a given site impacts the enrichment of mutations at a second site for (A) antigen binding (**Dataset #2**) and (B) non-specific binding (**Dataset #3**). For example, fixing residue T(H55) strongly impacts the enrichment of mutations at position H50 for antigen binding. Much less contextual dependence was observed for non-specific binding relative to antigen binding.

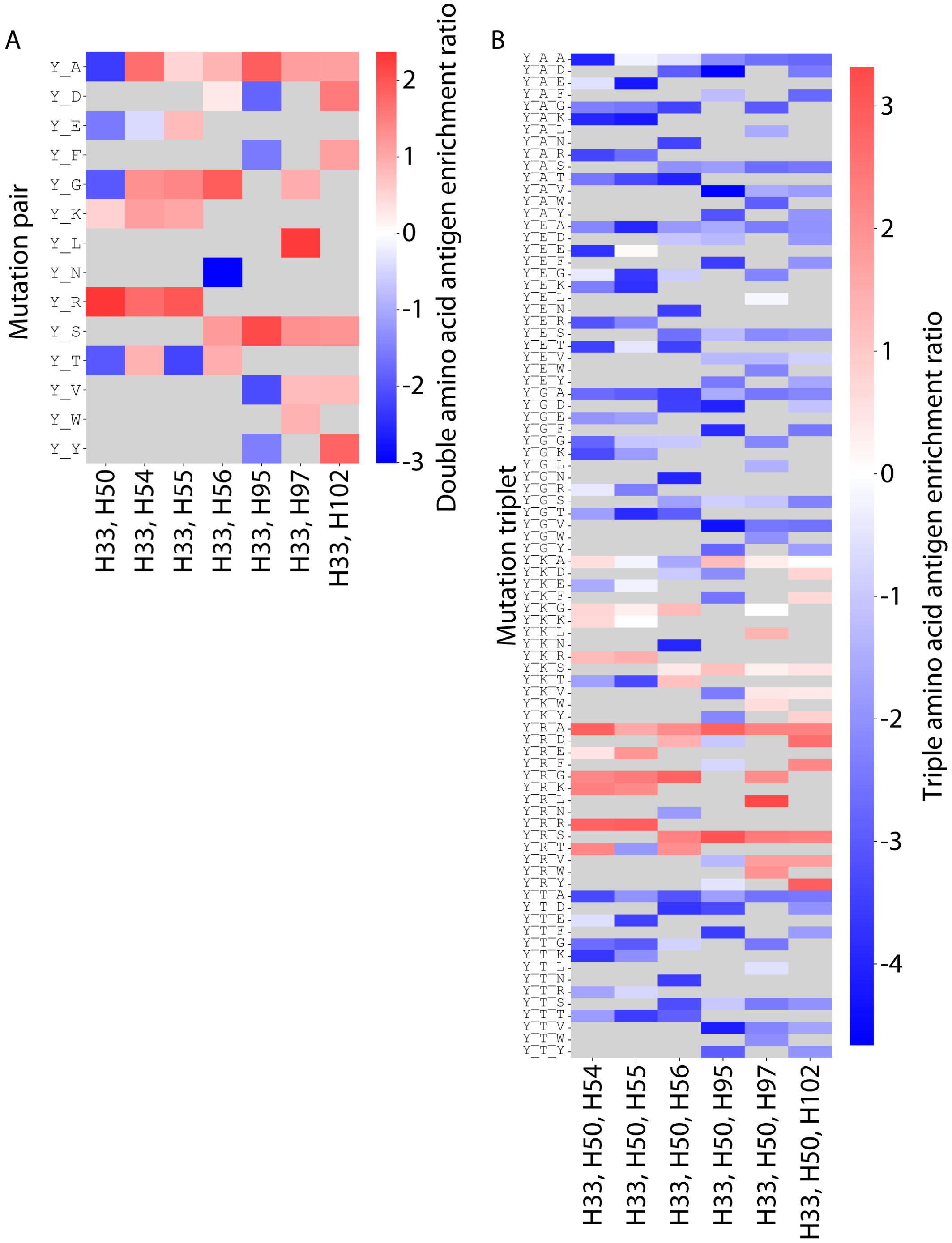


**Figure S7. Multi-PSERMs for antigen binding.** PSERMs were developed to score co-enrichment of sets of two residues (2-PSERM) or three residues (3-PSERM), and a subset of results are shown. (A) 2-PSERM is shown in which residue Y(H33) is paired with every other sampled residue at the other positions. (B) 3-PSERM is shown in which residue Y(H33) is paired with every other pair of residues that includes position H50 as well as the other six mutated positions. Each value of these matrices represents the enrichment of pairs or triplets of residues. Red denotes enrichment and blue denotes depletion.
